# Interconnections accelerate collapse in a socio-ecological metapopulation

**DOI:** 10.1101/195412

**Authors:** Zachary Dockstader, Chris T. Bauch, Madhur Anand

## Abstract

Resource over-exploitation can have profound effects on both ecosystems and the human populations residing in them. Models of population growth based on a depletable resources have been studied previously, but relatively few consider metapopulation effects. Here we analyze a socio-ecological metapopulation model where resources grow logistically on each patch. Each population harvests resources on its own patch to support population growth, but can also harvest resources from other patches when their own patch resources become scarce. We find that allowing populations to harvest from other patches significantly accelerates collapse and also increases the parameter regime for which collapse occurs, compared to a model where populations are not able to harvest resources from other patches. As the number of patches in the metapopulation increases, collapse is more sudden, more severe, and occurs sooner. These effects also persist under scenarios of asymmetry and inequality between patches. We conclude that metapopulation effects in socio-ecological systems can be both helpful and harmful and therefore require urgent study.

## 1. Background

Simple population dynamic models have long been used in theoretical population biology, beginning with the logistic growth model developed by Verhulst [1]:

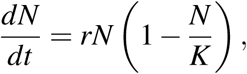

where *N*(*t*) is the population size at time *t*, *r* is the net growth rate, and *K* is the carrying capacity. This model represents resource-limited population growth reaching a carrying capacity *K* that is the largest population size that the resources of the environment can support. The logistic growth model and various extensions thereof are richly represented in the ecological literature and have been used as a framework to study population dynamics in a variety of species [2, 3, 4].

Other literature has extended single population models to study metapopulation dynamics. A metapopulation is a collection of spatially distributed populations all belonging to the same species [5]. Metapopulation models provide important insight into interactions of connected populations. Previous research on metapopulations has identified phenomena such as the rescue effect and extinction debt. In a rescue effect, the local extinction of a population is prevented due to immigration of a neighboring population residing in the same metapopulation [6]. In contrast, extinction debt refers to an effect whereby destruction of a natural habitat has not only immediate impacts on populations, but also creates a ‘debt’ effect whereby future extinctions will occur as well long after habitat destruction has ceased [7].

Although population growth models such as the logistic growth model have arguably found their fullest expression in ecology, Verhulst developed his model for application to human populations and he inferred the model’s parameter values using population data from Belgium and other countries. This interest in human populations may have been due to the influence of Thomas Malthus and his work ‘An Essay on the Principle of Population’, which is well known for its hypothesis that famine and poverty are mathematically inevitable [8].

Malthus continues to influence our thought in a time of severe global over-consumption and resource depletion. Resource depletion has been conceptualized and quantified in various ways. For instance, recent literature identifies seven planetary boundaries that must not be transgressed if humans are able to live sustainably on the planet, and finds that three of these boundaries have already been transgressed [9].

More recently, researchers have introduced frameworks to describe the process by which over-consumption occurs. For instance, the red/green sustainability framework describes how populations become increasingly disconnected from their impacts as they urbanize [10]. In a ‘green’ phase, populations are highly dependent on their local environment for their subsistence, and therefore feedback from environmental implications of human activity are quick to down-regulate the human activity. However, as populations develop and draw their resources from a global resource pool, their economic activities cause environmental impacts that are no longer felt by them but rather by geographically distant populations, weakening the short-term coupling between humans and their environment. This process is captured by, for instance, the linkages between local deforestation and high pressure for international agricultural exports [11], and the large dependence seafood markets in Japan, the United States, and the European Union on foreign sources [12].

Although logistic growth models and its variations are most widely used in ecology, the application of population growth models to resource-limited human populations has also received attention, perhaps on account of our growing awareness of the possible ramifications of resource over-exploitation, especially in the face of environmental change [13]. Mathematical models have been used to study phenomena such as human population collapse in a model with resource dynamics [14] and conflict among metapopulations arising over common resources [15, 16]. Models have also been used to study historical human population collapses such as in the people of Easter Island [17, 18, 19, 20, 21, 22, 23, 24, 25, 26, 27], the Kayenta Anasazi [28], and the Andean Tiwanaku civilization [29], as well as collapse of modern populations [30, 31, 32, 33].

With some exceptions [15, 16, 27, 34], most previous research on resource-limited population growth focuses on single populations and not on multi-population interactions. However, multi-population interactions through trade and other mechanisms are an inescapable feature of the world’s human metapopulation dynamic, and can have significant impacts on ecosystems and resource levels [35]. The literature on multi-population interactions has explored meta-population models involving migration of individuals between patches [28], or competing populations conflicting and bargaining over a common resource [15, 16]. A multi-patch model of human populations fitted to data from Easter Island has predicted that coupling human populations together through exchange of resources, migration and technology can stabilize the entire metapopulation [27].

In this paper we build on a previous single population model [18] to create a metapopulation model of resource-limited growth that captures mechanisms similar to the red/green sustainability framework. Populations grow logistically by exploiting a depletable resource that obeys a resource dynamic. Local populations can take resources not only from their own patch but also from other patches, when resources in their own patch become scarce. Our research objective was to determine whether the metapopulation collapses faster or more often when patches are allowed to harvest resources from other patches. We describe the model in the next section, followed by Results and a Discussion section.

## 2. Model

We build on a previous single-population model analyzed by Basener and Ross [18] who formulated a model whereby the population grows logistically to a carrying capacity that is proportional to the resource level. A second equation describes the logistic growth of resources to a separate carrying capacity, minus harvesting. We develop both two-patch and ten-patch versions of our model.

### Two-patch model

In the two-patch model, patch 1 has population size *P*_1_ and resource level *R*_1_, and patch 2 has population size *P*_2_ and resource level *R*_2_:

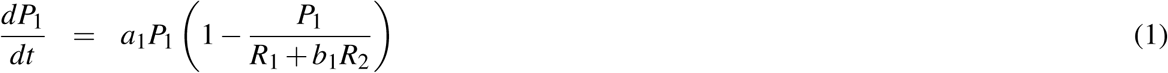

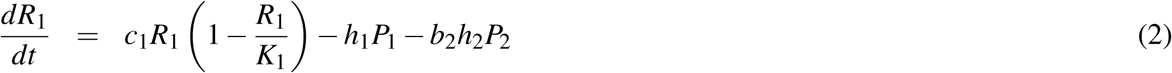

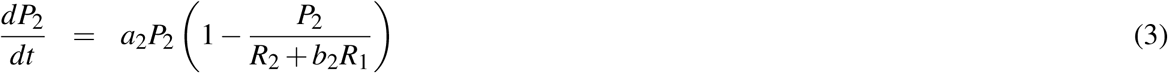

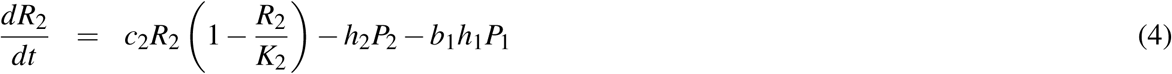

where *a*_1,2_ is the growth rate of patch 1 (resp. 2); *c*_1,2_ is the resource renewal rate in patch 1 (resp. 2); *K*_1,2_ is the carrying capacity of the depletable resource in patch 1 (resp. 2); *h*_1,2_ is the baseline harvesting rate at which patch 1 (resp. patch 2) harvests resources for its population’s consumption; *b*_1_ is the proportionof resources that patch 1 takes from patch 2, and similarly for *b*_2_. In this model, the carrying capacity of the human populations is determined by how much resource is available to support them, either from their own patch or taken from the other patch. When *b*_1_ = *b*_2_ = 0 we recover the original model by Basener and Ross [18].

We set *b*_1_ = *b*_1_(*R*_1_*, P*_1_) and assume that patch 1 will attempt to harvest more resources from patch 2 when the resources from patch 1 are not enough to support the patch 1 population. Similarly, *b*_2_ = *b*_2_(*R*_2_*, P*_2_). These functions take the form

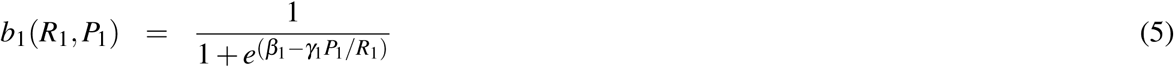

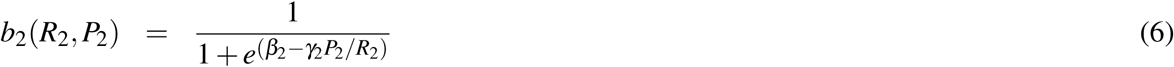

These are sigmoidal functions where the rate at which patch 1 harvests from patch 2 is higher when *P*_1_*/R*_1_ is higher, and *vice versa*, where *β*_1_ *>* 0 controls the location of the mid-point of the sigmoid, and where *γ*_1_ *>* 0 controls how steep the curve is. Parameters *β*_2_ and *γ*_2_ are similarly defined.

### Ten-patch model

We also analyzed a version of the model where ten patches are interconnected and can take resources from one another. The dynamics of patch *i* in the ten-patch model are given by

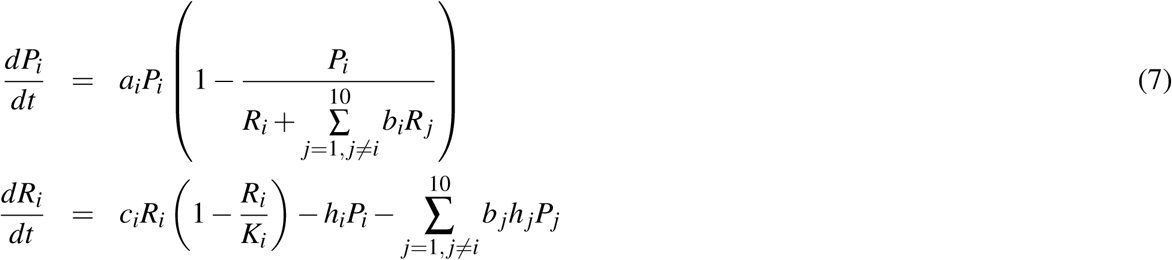

where parameter definitions are the same as in the two-patch case.

### Baseline parameter values

The baseline values of our parameters appear in Table 1. The population growth rate *a*_1,2_ was estimated by historical world population growth rates [36]. The resource growth rate *c*_1,2_ was taken as the average increase in crop yield since 1961 [37]. The values of the harvesting efficiency *h*_1,2_ and carrying capacity of the resources *K*_1,2_ were calibrated so that the populations would begin with enough resources to survive for several centuries regardless of their rates of resource use, and so their harvesting efficiency was high enough that there were consequences to over-exploitation but not high enough to make resource use incredibly costly. At these parameter values, the population size of a single patch grows to 650, 000 and then declines somewhat to an equilibrium population size of 480, 000 over a timescale of several hundred years.

**Table 1.** Baseline model parameter values.

| Symbol | Definition | Value | Source |
| --- | --- | --- | --- |
| $a_{1,2}$ | Population 1,2 net growth rate | 0.0177/year | [36] |
| $c_{1,2}$ | Resource growth rate in patch 1, 2 | 0.015/year | [37] |
| $h_{1,2}$ | Harvesting efficiency of population 1, 2 | 0.008/year | calibrated |
| $K_{1,2}$ | Carrying capacity of resources in patch 1, 2 | 1,000,000 | calibrated |
| $\beta_{1,2}$ | Controls location of the mid-point of the sigmoid for population 1, 2 | 3.5 | calibrated |
| $\gamma_{1,2}$ | Controls steepness of the sigmoid for population 1, 2 | 5 | calibrated |

The parameters controlling the midpoint and steepness of the sigmoid function (*β* and *γ*) were obtained through calibration by analyzing the effect they had on how much and when the populations would take from neighboring populations. To calibrate *β*, our intention was that the populations did not take much from neighbors when they were not in need. In contrast, they would take more when their resources began to dwindle and neighbour’s resources were needed to survive. To calibrate *γ* we choose a value such that the switch between these two described states was relatively gradual. In particular, we required *β_i_* and *γ_i_* to satisfy the property that if *P*_*i*_/*R*_*i*_ = 1*/*2 and thus resources were abundant, then *b*_*i*_ was roughly 25%, whereas if *P*_*i*_/*R*_*i*_ = 1, indicating a situation where a shortage of resources was beginning to become worrisome, then *b*_*i*_ would be greater than 75%.

Initial conditions were *P*_1_(0) = 50, 000, *P*_2_(0) = 50, 000, *R*_1_(0) = 1, 000, 000, and *R*_2_(0) = 1, 000, 000. These initial conditions corresponded to two populations with relatively low starting population levels and with initially abundant resources at carrying capacity in their respective patches.

We solved the model equations numerically using the adaptive fourth-fifth order Runge Kutta method implemented via Matlab’s ODE45 solver. The code can be found on Github [38]. We compared model dynamics for both interconnected and isolated versions of the two-patch and ten-patch models to determine the impact of interconnectedness on the likelihood and timing of collapse.

We explored the sensitivity of model predictions to parameter variations away from the baseline parameter values. In this process we considered two different scenarios for parameter variations. In most figures, populations were always assumed symmetric and had identical parameter values, but some figures explore the case of asymmetric populations where the two populations differ in one parameter value.

## 3. Results

### 3.1. Baseline Scenario

The baseline scenario was simulated for both the interconnected (*b*_1_*, b*_2_ *>* 0) and isolated (*b*_1_ = *b*_2_ = 0) versions of the model. In the interconnected baseline scenario (Fig. 1), the populations begin a nearly exponential increase in their population growth (Fig. 1a) as they quickly reduce their local resources (Fig. 1b). This decrease in local resources causes the populations to begin taking resources from their neighboring patch to continue supporting their population (Fig. 1d). This results in greater resource availability (Fig. 1c) which stimulates further unsustainable population growth. Once the resources of both patches are strongly depleted, both populations collapse.

**Figure 1.**
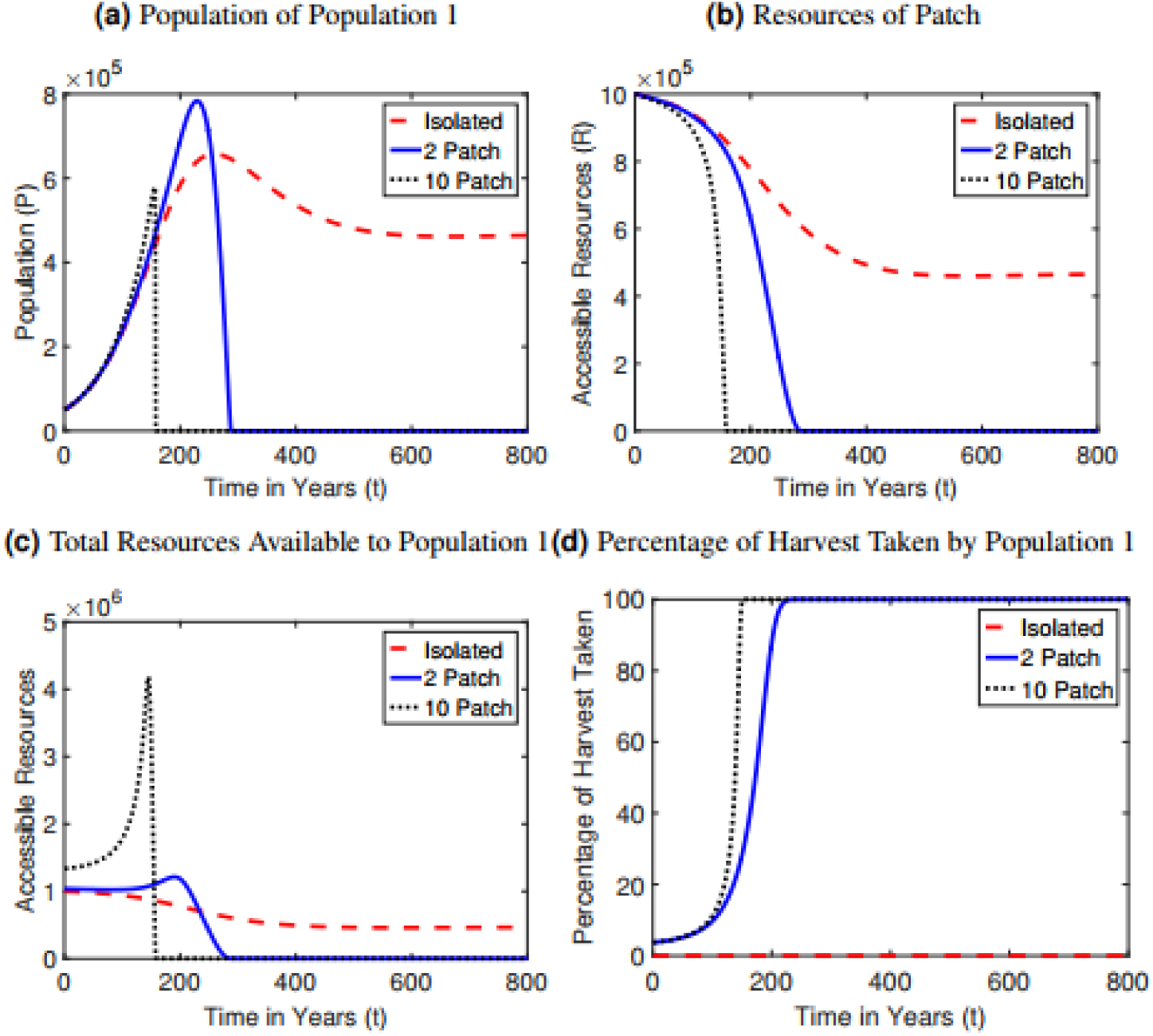
Model dynamics at baseline parameter values for the isolated, interconnected, and 10 population scenarios for (a) population size *P*_1_ of patch 1, (b) resource availability *R*_1_ in patch 1, (c) total resources available to population 1, and (d) percentage of harvest of patch 2 taken by population 1. Results for population 2 and patch 2 are symmetrical.

In contrast to the interconnected case, both populations achieve sustainability in the isolated case at baseline parameter values (Fig. 1). Much like the interconnected scenario, the population of both civilizations grow very quickly, reaching a peak and then beginning to decline (Fig. 1a). However, instead of complete extinction of the population, the population decline begins to slow as the system reaches a steady state in which the population and resources equilibrate at an intermediate level, achieving sustainability.

Dynamics in the 10-patch model amplify the trends in dynamics observed in the 2-patch model. The initial increase in population size is much more rapid, but the following collapse happens much sooner and is much more sudden than in the 2-patch model (Fig. 1a). Collapse occurs after 159 years in the 10-patch model compared to the 289 years in the 2-patch model. Resources are depleted much more rapidly in the 10-patch model (Fig. 1b-d).

### 3.2. Time to collapse

We also studied how the time to collapse depends on parameter values for the isolated and interconnected scenarios. Time to collapse was defined as the time elapsed until the populations of both patches reaches zero (*P*_1,2_ *<* 10^-7^). We generated plots showing the time to collapse versus a single parameter, with all other parameter values held constant at their baseline values (Fig. 2,3). By doing so, we obtain an idea of whether the more rapid collapse of interconnected systems compared to isolated systems is robust to changes in model parameter values, and which parameter values are most influential in determining collapse.

**Figure 2.**
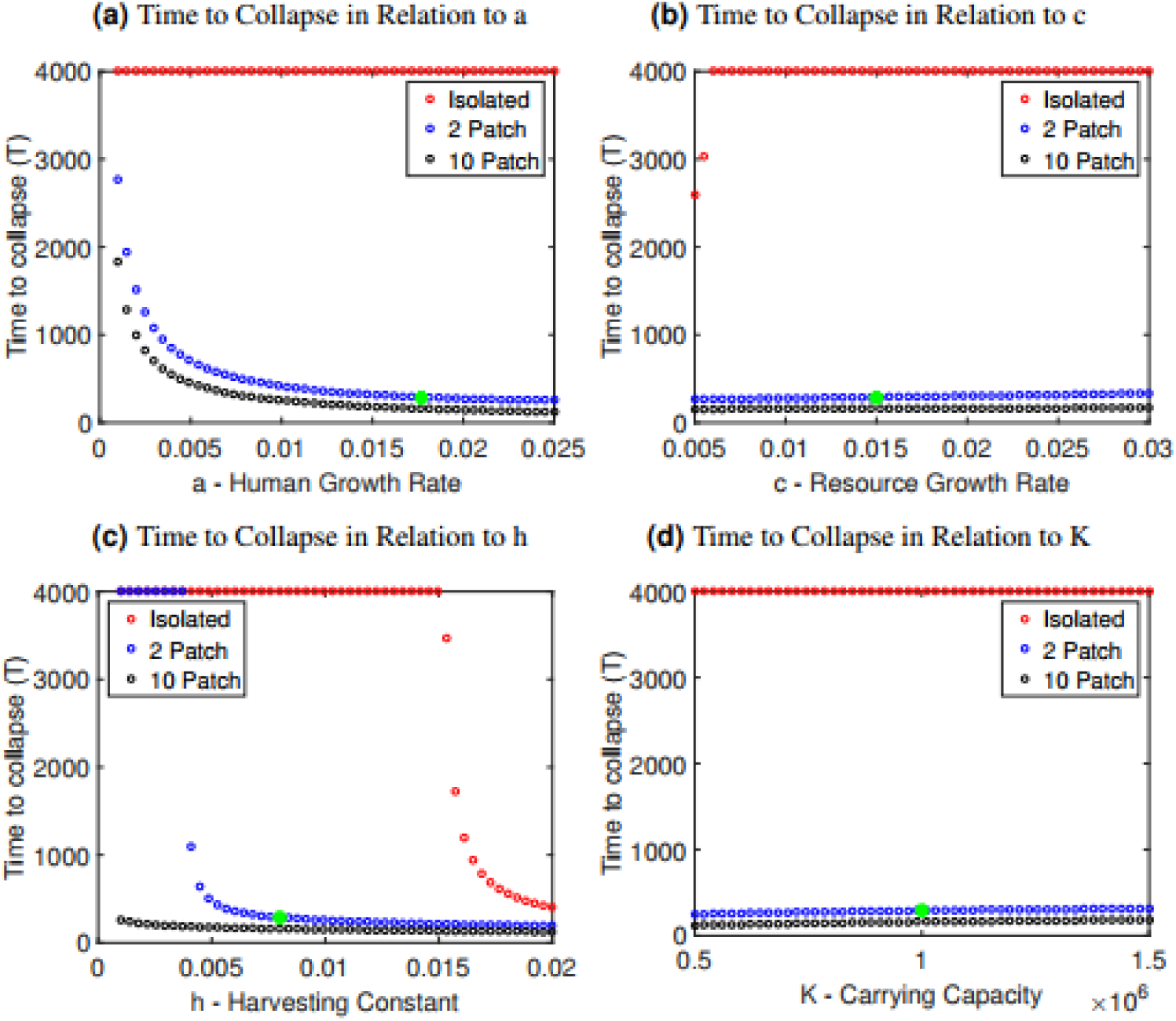
Time to collapse for the isolated and interconnected cases as it depends on changes in (a) the human growth rate *a*, (b) the resource growth rate *c*, (c) the harvesting constant *h* and (d) the carrying capacity *K*. The parameter along the horizontal axes was changed for both patches, thus preserving symmetry. A green star has been included in each graph to indicate the value of the parameter in the baseline scenario.

**Figure 3.**
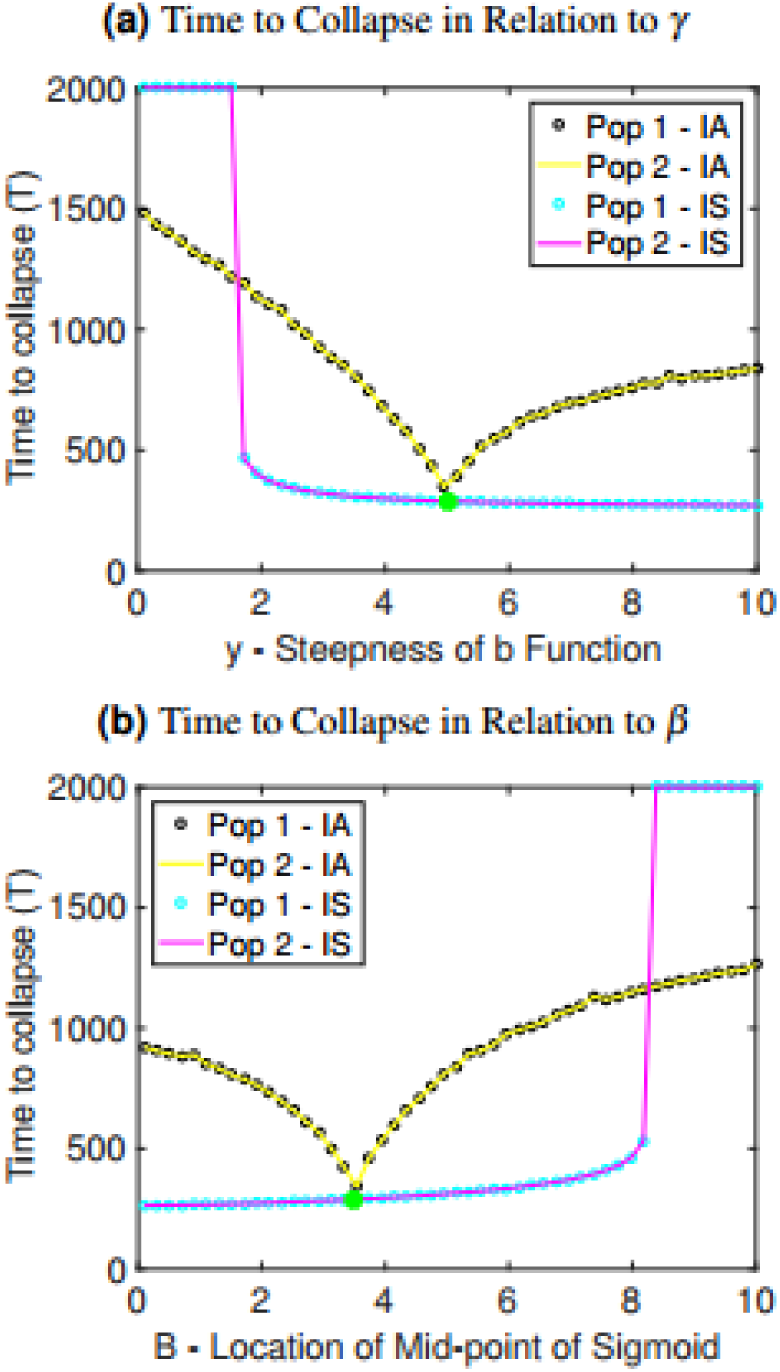
Time to collapse for the isolated and interconnected cases as it depends on changes in (a) the steepness of sigmoidal function *γ*, (b) the midpoint of sigmoidal function *β*. A green and yellow star have been included in each graph to indicate the value of the parameter in the baseline scenario of the interconnected case and isolated case, respectively. IS denotes interconnected symmetric, wherein the parameter along the horizontal axes was changed for both patches, thus preserving symmetry, while IA denotes interconnected asymmetric, wherein the parameter values for population 1 was changed while the parameter values for population 2 was held constant at its baseline value.

Across a broad range of parameter values, time to collapse in the interconnected case is much shorter, demonstrating that the interconnection of the two populations is detrimental to the stability of the system (Fig. 2, 3). The isolated case is more resilient to collapse, as we see that the model often survives indefinitely in all cases except when the harvesting constant or resource growth rate are changed drastically relative to the baseline values (Fig. 2b, c). This is in contrast to the interconnected case, where nearly all parameter choices for the human growth rate *a*, the resource growth rate *c*, the harvesting constant *h* and the carrying capacity *K* lead to collapse. Collapse occurs more rapidly when the human growth rate *a* or the harvesting constant *h* are increased, since both scenarios correspond to populations growing unsustainably quickly (*a*) or exploiting their resources unsustainably quickly (*h*). Interestingly, it is relatively independent of the carrying capacity *K* and the resource growth rate *c*. Therefore in this system, increasing carrying capacity (*K*) by boosting yield by, or increasing the ability of the resource to replenish itself (*r*) has relatively little effect in delaying the collapse.

The more rapid collapse observed in the 10-patch model compared to the 2-patch model (Fig. 1) is also robust under these parameter variations (Fig. 2). As the number of populations increases from 2 to 10, the time to collapse declines with the number of populations (Fig. 4).

**Figure 4.**
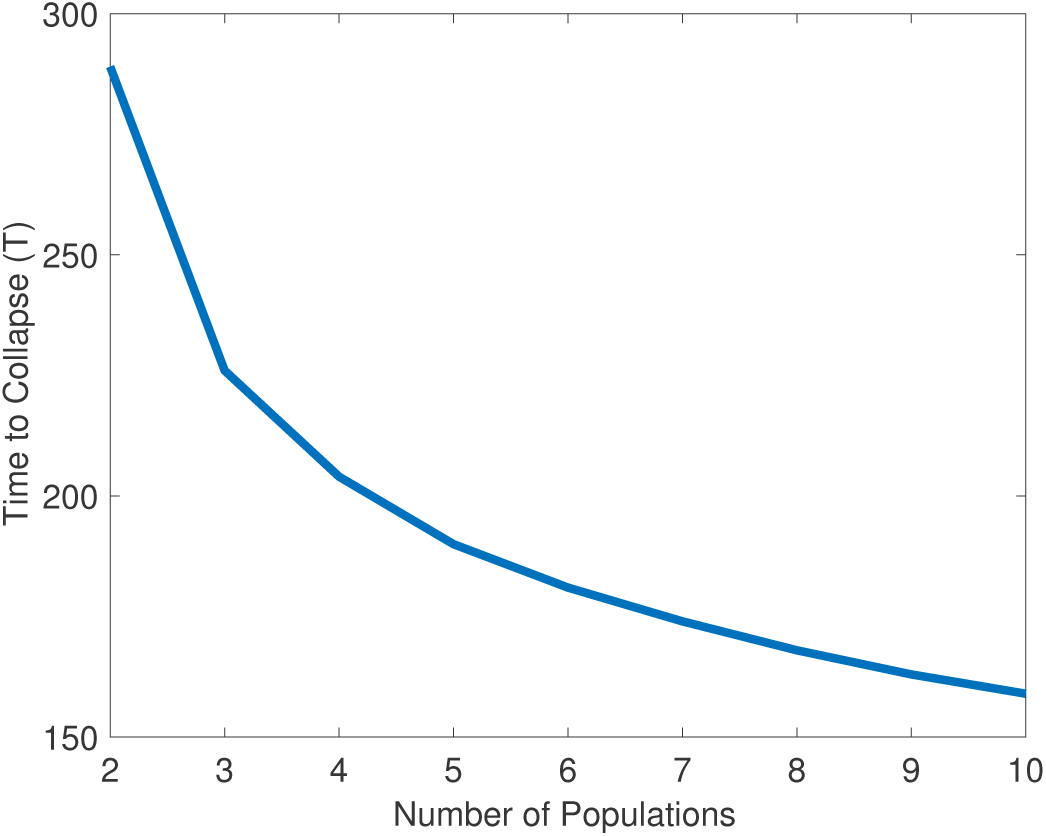
Time to collapse versus number of populations included in model. Baseline parameter values were used (Table 1).

The observed relationships between time to collapse and interconnectedness are also preserved under variation in parameters controlling the rate at which one patch harvests resources from another patch: *β*, which controls the midpoint location in the sigmoidal function, and *γ*, which controls the steepness of the sigmoidal function (Fig. 3). When *γ* is increased, the switch to harvesting from other patches happens more quickly, causing more rapid collapse (Fig. 3a). Interestingly, if *γ* is sufficiently low (meaning the sigmoidal function transitions smoothly), then collapse does not occur. Hence, if populations transition more gradually to harvesting from other patches, collapse can be avoided. When *β* is decreased, populations begin harvesting from other patches earlier and more intensely, causing more rapid collapse (Fig. 3b).

The case of asymmetric parameter variation is also considered in Fig. 3 to provide a contrast with our baseline assumption of symmetric parameter values. As the value of *γ* is increased for only one of the populations while the value of *γ* for the other population is held constant, the time to collapse decreases for both populations until it reaches a minimum at the baseline value, and then starts to increase again (Fig. 3a). Similarly, if *β* is increased for only one of the populations, time to collapse decreases until it reaches the baseline value but then increases again (Fig. 3b). This suggests that heterogeneity in the metapopulation may stave off collapse.

### 3.3. Parameter planes

By varying two parameters at one time and holding all others constant at their baseline values, we can understand parameter combinations that lead to collapse or survival under the isolated and interconnected scenarios. It is evident from these parameter planes that the isolated case of the model is far less prone to collapse over the same ranges of parameter values. Collapse occurs for a much wider part of the parameter plane under the interconnected symmetric case than under the isolated case (electronic supplementary material, Figure S1). In contrast to the baseline parameter values, we observe parameter regimes in the interconnected symmetric case where increasing the resource growth rate *c* can move the populations into a region of sustainability. Introducing asymmetry to the parameter plans, such that the two parameter values for one population are varied while the parameter values for the other population are held at baseline values, we observe that sustainability is a more frequent outcome than in the symmetric case, but occurs less frequently than in the isolated case (electronic supplementary material, Figure S1).

### 3.4. Impact of Inequality

To observe the effect of inequality on system dynamics, we created an additional scenario involving two unequal populations. Population 1 has a higher starting population size, population growth rate, resource growth rate and harvesting efficiency, but a lower carrying capacity than population 2, which has more resources but a lower starting population size and growth rate. Population 1 is also more prone to take resources from population 2 than *vice versa*. The inequality scenario was simulated with and without interconnections. Parameter values can be found in electronic supplementary material, Table S1 and the initial conditions were *P*_1_(0)=50,000, *P*_2_(0)=25,000, *R*_1_(0) = 250,000, *R*_2_(0)=1,000,000.

In the interconnected case (electronic supplementary material, Figure S2), population 1 grows relatively quickly (Fig. S2a), reaching their maximum population size nearly 100 years before population 2. In the process, they exhaust all of their resources early in the simulation (Fig. S2b). However, this causes very little disturbance to population 1 since there is only a small, nearly non-existent, decrease in population size at the time of resource depletion. This is due to their early dependence on population 2’s resources (Fig. S2g) dampening the effect that over-exploitation has on their own population. After this point, both populations continue to consume population 2’s resources (Fig. S2d) until the inevitable depletion, causing both populations to collapse.

In the corresponding isolated but unequal case (electronic supplementary material, Figure S3), the outcomes are very different. Population 2 begins a similar population increase as in the interconnected case, but the population avoids complete collapse and instead recovers to a stable state (Figure S3c). However, population 1 grows unsustainably, over-depletes their resource, and collapses (Fig. S3a,b). Hence, for these parameter values, we observe that the dichotomy between outcomes in the isolated and interconnected scenario persists when the two populations are unequal.

## 4. Discussion

In this paper we extended a simple population model where a population harvests a depletable resource to a metapopulation setting where a population patch can also harvest resources from other patches when their own resources run low. We showed how the populations collapse faster and for a broader range of parameter values when patches are allowed to harvest resources from other patches. As the number of patches increases, the effect is amplified.

Interconnections accelerate collapse in this model because the ability to harvest resources from other patches enables populations to access a larger resource pool. Consequently, the populations are able to grow at a very rapid rate, compared to the case where patches are isolated from one another. Each patch population size grows beyond what is sustainable using only the resources in a single patch, and this causes rapid collapse as the resources disappear and all patches are left with unsustainably high populations. This mechanism operates even when the net resource growth rate *c*_1,2_ parameter exceeds the net population growth rate parameter *a*_1,2_. Collapse remains possible in the isolated scenario, but the smaller available resource pools tend to prevent it.

This effect was robust under a wide range of parameter variation. We also found that asymmetry in parameter values between the two patches does not change the qualitative results, but does tend to stave off collapse. We speculate that models with greater heterogeneity (such that each patch has a unique set of parameter values) might replicate this feature, but we leave this for future work. We furthermore found that collapse can occur in a scenario of inequality between the two patches, although we did not test the robustness of this finding to parameter variation.

In some respects, our model embodies some of the ideas of “red and green loop” dynamics as introduced by Cumming et al [10]. Populations in our model can depend on resources harvested non-locally, such that the population is buffered from the implications of their harvesting activities in the short term. Thus, in the interconnected case, populations are in a red loop regime. As the population transitions to relying on the resources of other patches as its own resources are depleted, the red loop progresses to a red trap corresponding to collapse of both populations in the interconnected scenario. In comparison, in the isolated case, populations are much more dependent on their local resources and feel the impacts of their harvesting choices immediately: a green loop regime.

Our model makes simplifying assumptions that may influence its predictions. For instance, due to the structure of our sigmoidal function governing cross-patch harvesting and in particular the assumed dependence of cross-patch harvesting on *P*_*i*_/*R*_*i*_, patches tend to collapse simultaneously when **R*_*i*_* becomes small. Moreover, patches cannot prevent cross-patch harvesting. In reality, effective institutions (where they exist) would be able to prevent cross-patch harvesting through legislation and this might have the effect of preventing collapse from spreading to all patches. Future work could study the effects of retaining a portion of local resources for the native patch’s exclusive use. Similarly, allowing migration of individuals as well as cross-patch harvesting could influence dynamics, perhaps even to the point of preventing collapse [27]. Non-human species migrate when local resources are depleted; humans migrate but technology now allows them to import the resources they need without migrating. Allowing cross-patch harvesting while preventing migration could therefore be particularly dangerous.

Similarly, we assumed a Malthusean world where more resources are always converted into more offspring. However, it is observed that most populations go through a demographic transition to lower fertility when they become sufficiently industrialized [39]. Incorporating this effect into the model may help prevent unsustainable growth, although the strength of the effect depends on whether increases in per capita resource consumption outstrip the benefits of slowed population growth.

Another possible extension of the model is to include dynamically changing parameters. At the moment, all parameters in the model are static. However, technological improvements mean that parameters like the harvesting efficiency *h* and cross-patch harvesting should change over the course of the simulation. In this vein, work by Reuveny and Decker [25] explores how technological advancement affects a human-resource population model. Similarly modifications to our model could be implemented, and their effects studied. In our multi-population socio-ecological model where populations grow by harvesting a depletable resource, the ability of one patch to support its population growth by harvesting resources from other patches increases population growth in the short run, but causes population collapse in all patches in the long run. This effect is robust to parameter variation, and is accelerated significantly by the inclusion of more patches. Given the ubiquity of cross-patch harvesting in real populations, socio-ecological models of human growth and resource consumption should consider the role of metapopulation effects.

## Competing Interests

The authors declare no competing interests.

## Author Contributions

M.A. and C.T.B. conceived the study; Z.D. analyzed the model and generated figures; all authors wrote and reviewed the manuscript.

## Acknowledgements

This research was supported by NSERC Discovery Grants to C.T.B. and M.A.

